# Cyanotoxins accumulate in Lake St. Clair fish yet their fillets are safe to eat

**DOI:** 10.1101/2022.09.08.507173

**Authors:** René S. Shahmohamadloo, Satyendra P. Bhavsar, Xavier Ortiz Almirall, Stephen A. C. Marklevitz, Seth M. Rudman, Paul K. Sibley

## Abstract

Consuming fish exposed to cyanobacterial harmful algal blooms (HABs) may be a major route of microcystin toxin exposure to humans. However, it remains unknown whether fish can accumulate and retain microcystins temporally in waterbodies with recurring seasonal HABs, particularly before and after a HAB event when fishing is active. We conducted a field study on Largemouth Bass, Northern Pike, Smallmouth Bass, Rock Bass, Walleye, White Bass, and Yellow Perch to assess the human health risks to microcystin toxicity via fish consumption. We collected 124 fish in 2016 and 2018 from Lake St. Clair, a large freshwater ecosystem in the North American Great Lakes that is actively fished pre- and post-HAB periods. Muscles were analyzed using the MMPB Lemieux Oxidation method for total microcystins, which was used to perform a human health risk assessment for comparison against fish consumption advisory benchmarks available for Lake St. Clair. From this collection 35 fish livers were additionally extracted to confirm the presence of microcystins. Microcystins were detected in all livers at widely varying concentrations (1-1,500 ng g^-1^ ww), suggesting HABs are an underappreciated and pervasive stressor to fish populations. Conversely, microcystin levels were consistently low in muscles (0-15 ng g^-1^ ww) and presented negligible risk, empirically supporting that fillets may be safely consumed before and after HAB events following fish consumption advisories.

## 1. Introduction

The global expansion of cyanobacterial harmful algal blooms (HABs) is a pressing challenge for freshwater ecosystems. Their spread is due, in part, to climatic changes and an expansion in human activities, particularly intensified agriculture, which render waterbodies increasingly vulnerable to biodiversity loss and threatened ecosystem services (Hellweger et al., 2022; Ho et al., 2019; Huisman et al., 2018; Reid et al., 2019; Smith et al., 2019). Freshwater HABs can produce microcystins, potent hepatotoxins that primarily inhibit protein phosphatases and at sufficient levels can cause liver necrosis, tumor formation, and organ failure in humans and wildlife (Chorus and Welker, 2021; Shahmohamadloo et al., 2023b, 2022, 2021).

The World Health Organization’s (WHO) guidelines (WHO, 2020) are implemented in many countries to protect humans from microcystin toxicity in drinking water (12 µg L^-1^ short-term, 1 µg L^-1^ lifetime); however, longstanding concerns remain that fish consumption may be a viable route of algal toxin exposure (FAO, 2020; IPCC, 2022). These concerns stem from the possibility that microcystins may be one of several factors involved in fish kills during HAB events (Ibelings et al., 2005) or that microcystins can accumulate up the food chain (Gene et al., 2019; Shahmohamadloo et al., 2020a, 2020b). Previous studies measured microcystins in fish tissues at levels exceeding the WHO’s tolerable daily intake (TDI) value (0.04 µg kg^-1^ body weight per day) using the enzyme linked immunosorbent assay (ELISA) (Dyble et al., 2011; Poste et al., 2011; Wituszynski et al., 2017). Yet, ELISA is prone to false positives in fish tissues because it only measures free fractions of microcystins, whereas bound fractions can make up to 90% of the total amount of toxins present in a sample (Birbeck et al., 2019; Flores et al., 2018; Schmidt et al., 2014). This has raised skepticism about the accuracy of such data in assessing risks to humans via consumption of fish. Intertwined in the debate about methodology are projections stating that climate change will increase the mean number of days of a HAB event from 16-23 days in 2050 to 18-39 days in 2090 in North America (Chapra et al., 2017).

Recent work in Lake Erie addresses some of the methodological and ecological concerns by using an optimized MMPB Lemieux Oxidation method (Anaraki et al., 2020) to dually assess the human health risks from fish consumption and fish health risks from microcystin toxin exposure before, during, and after HAB events (Shahmohamadloo et al., 2023a). Interestingly, the findings suggest both cause for relief and concern: while Lake Erie fish fillets may be safe to eat, fish livers can retain high levels of microcystins (mean 460.13 ng g^-1^ww across species) and may be a pervasive threat to fish populations (Shahmohamadloo et al., 2023a). Lake St. Clair, a large waterbody that is important for recreational fishers (OMNRF, 2016), regularly experiences microcystin-HABs (Davis et al., 2014) and may expose resident fish populations in the future to blooms with extended residency, severity, and toxicity. Aware that the present duration of a HAB (Chapra et al., 2017) is a small fraction of the total fish season and the overwhelming body of literature on fish was collected during HABs (Dyble et al., 2011; Poste et al., 2011; Wituszynski et al., 2017), there is an invested interest to determine the human health risks from consuming fish before and after HAB events when recreational fishing is more active.

We collected 124 recreationally important sport fish, including some species which have received little attention previously, namely Largemouth Bass (*Micropterus salmoides*), Northern Pike (*Esox lucius*), Smallmouth Bass (*Micropterus dolomieu*), Rock Bass (*Ambloplites rupestris*), Walleye (*Sander vitreus*), White Bass (*Morone chrysops*), and Yellow Perch (*Perca flavescens*) from Lake St. Clair in 2016 and 2018 throughout pre- and post-HAB periods when recreational fishing is more active. We measured microcystins in fish muscles using the MMPB Lemieux Oxidation method (Anaraki et al., 2020). In addition, we dissected 35 livers from these samples to confirm the presence of microcystins in fish tissues and assess the amount retained pre- and post-HAB periods. Results from muscles were then compared against fish consumption advisory benchmarks for microcystins available to Lake St. Clair and applied in a human health risk assessment to determine whether fish tissues exceeded the WHO’s TDI. Here, we hypothesize that microcystin levels in fish muscles will vary among fish species but, overall, will pose low human health risks in pre- and post-HAB periods.

## 2. Methods

### 2.1. Study location and water sampling

Eight locations in Lake St. Clair (42° 25’ 20” N, 82° 39’ 36” W) were sampled for fish in 2016 and 2018 during pre- (June to July) and post- (October to November) HAB periods. During this time, *Microcystis, Planktothrix*, and *Dolichospermum* are typically present in Lake St. Clair, with toxigenic strains of *Microcystis* predominating (Davis et al., 2014). Physical, chemical, and biological water quality monitoring HABs data was collected by the Ontario Ministry of Natural Resources and Forestry (MNRF; see Dataset S1). Water data was collected during repeated bi-weekly sampling trips ‘pre’, ‘during’, and ‘post’ HAB periods to characterize algae blooms and potential microcystin toxins. This corroborates recent findings demonstrating chlorophyll *a* is a consistent correlate of microcystins in freshwaters and can better account for the inevitable temporal variation associated with changing causes of microcystin production (Qian et al., 2021). Notwithstanding, the Ontario Ministry of the Environment, Conservation and Parks (MECP) additionally collected water samples during pre- and post-HAB periods from the southern shoreline of Lake St Clair near Belle River (42° 18’ 11” N, 82° 42’ 46.7” W) and Puce River (42° 18’ 4” N, 82° 42’ 19” W) to measure total phosphorus, chlorophyll *a*, and total microcystins. Total phosphorus (28.7 and 13.4 μg L^-1^), chlorophyll *a* (9.7 and 4.6 μg L^-1^) and total microcystins (0.2 and 1.0 μg L^-1^) were highest in 2018. Microcystin concentrations in the present study are similar to previous measurements from Lake St. Clair (Davis et al., 2014; Taranu et al., 2019).

### 2.2. Fish sampling

A total of 124 samples from Largemouth Bass, Northern Pike, Smallmouth Bass, Rock Bass, Walleye, White Bass, and Yellow Perch (Table 1, Dataset S1) were collected in 2016 and 2018 for microcystin screening as a part of the MECP’s Fish Contaminant Monitoring Program (Etobicoke, Ontario, Canada) in partnership with the MNRF (Wheatley, Ontario, Canada). Fish length and weight were recorded for all samples, after which the dorso-lateral muscle tissues (i.e., skinless, boneless fillets) were dissected for microcystin analysis. In 2018, a subset of 35 samples of Walleye, White Bass, and Yellow Perch were additionally dissected for livers to confirm their exposure to microcystins. All muscles and livers were frozen at -80 °C until homogenization and microcystin analysis.

**Table 1.** Fish collected from Lake St. Clair pre- and post-HAB for microcystin analysis in livers and muscles. Data is reported in mean±SD.

| Species | *n* | Length (cm) | Weight (g) | Microcystins in liver (ng g<sup>-1</sup> ww) | Microcystins in muscle (ng g<sup>-1</sup> ww) | EDI (μg kg<sup>-1</sup> body weight day<sup>-1</sup>)<sup>a</sup> |
| --- | --- | --- | --- | --- | --- | --- |
| Largemouth Bass | 3 | 38±13 | 1133±833 | - | 1.9±1.4 | 0±0.005 |
| Northern Pike | 12 | 74±10 | 2500±1055 | - | 0.4±0.9 | 0±0.003 |
| Rock Bass | 16 | 23±3 | 250±82 | - | 1.1±2.1 | 0±0.007 |
| Smallmouth Bass | 25 | 43±5 | 1334±559 | - | 1.0±3.1 | 0±0.01 |
| Walleye | 42 | 48±10 | 1059±690 | 217.2±220.5 | 1.8±2.5 | 0±0.008 |
| White Bass | 12 | 36±3 | 566±161 | 120.4±17.8 | 0.6±1.3 | 0±0.004 |
| Yellow Perch | 16 | 25±2 | 161±59 | 233.9±359.8 | 1.9±2.4 | 0±0.008 |

<sup>a</sup> Estimated daily intake (EDI) is measured as ng g<sup>-1</sup> d<sup>-1</sup>, assuming one fish meal is 227 g and applying the mean tissue concentration as the daily intake value. Values are compared against the World Health Organization's tolerable daily intake value (0.04 μg kg<sup>-1</sup> body weight day<sup>-1</sup>).

The total length of fish was measured because most resource users (i.e., fishers) and fish consumers are unable to determine the age of the fish they consume. For this reason, fish consumption advisories provided to the public, as well as health and resource managers are given based on fish length (MECP, 2017). By sampling and presenting the results based on length, we have provided real-world recommendations using metrics that the general population and decision-makers can access to help evaluate real-world risks and aid real-world decisions.

### 2.3. Fish tissue analysis

Fish tissues were analyzed for total microcystins (free and bound) using an optimized MMPB Lemieux Oxidation method that provides the necessary sensitivity and specificity (Anaraki et al., 2020)). Briefly, frozen tissues were freeze-dried using a Labconco Freezone 2.5 freeze-drier from Fisher Scientific (Markham, Ontario, Canada) for at least 24 h and then ground to a fine powder using a pestle and mortar. Next, approximately 100 mg of freeze-dried sample was spiked with the internal standard (*d*_*3*_-MMPB), oxidized at pH 8.5 using potassium permanganate (0.3 mM) and sodium periodate (20 mM) for 2 h. The oxidizing sample was quenched by adjusting the pH to 3 with 10% sulfuric acid and sodium bisulphite. Samples were then centrifuged for 7 min at 5,000 × g and correspondingly loaded onto Oasis HLB 3 cc (400 mg) LP extraction cartridges (Mississauga, Ontario, Canada) for clean-up and extraction of MMPB and *d*_*3*_-MMPB with 0.1% acetic acid and 50% methanol washes. Analytes were eluted with 4 mL of 100% methanol, dried down to 250 μL, and diluted to 1 mL using 0.1% FA milli-Q water. Samples were finally filtered using Pall GHP filters (Mississauga, Ontario, Canada) and prepared for instrumental analysis.

Samples were quantified using an *in-situ* generated MMPB matrix-matched calibration curve by isotope dilution with *d*_*3*_-MMPB by liquid chromatography coupled to time-of-flight mass spectrometry (LC-QTOF MS) using an Acquity I Class Chromatograph by Waters Corporation (Milford, Massachusetts, United States). This approach estimates potential matrix effects that may have impacted the derivatization, sample preparation and instrumental analysis steps. This method showed 16.7% precision (RSD) and +6.7% accuracy (bias), with a calculated method detection limit (MDL) of 2.18 ng g^-1^ wet weight (ww).

### 2.4. Human health risk assessment

Total microcystins measured in muscles were compared against Ontario’s fish consumption advisory benchmarks for microcystin-LR developed by the MECP and Ohio’s fish consumption advisory benchmarks for total microcystins developed by the Ohio Environmental Protection Agency. These benchmarks are based on the World Health Organization’s recommended tolerable daily intake (TDI) of 0.04 µg kg^-1^ bodyweight per day (WHO, 2020). The TDI value is based on the no-observed-adverse-effect-level (NOAEL) liver pathology observed in a 13-week study on mice using microcystin-LR (Fawell et al., 1994) and applying an uncertainty factor (UF) of 1000 (Eq. 1).

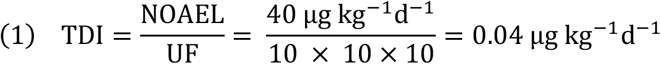

Ontario’s advisory benchmark assumes that 50% of microcystin exposure is through fish consumption. Microcystin-LR is one of the most toxic microcystin congeners (Shimizu et al., 2013). Therefore, application of a microcystin-LR advisory benchmark to total microcystins presents a more conservative and health protective scenario.

We further calculated the estimated daily intake (EDI) for total microcystins if one meal of fish is consumed every day (Eq. 2) and compared this to the WHO’s TDI for microcystin-LR.

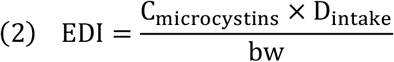

Where C_microcystins_ is the mean total concentration of microcystins in fish tissue wet weight (Dataset S1), D_intake_ is the daily fish consumption for Ontarians (one meal is 227 g) following the Ontario’s Guide to Eating Ontario Fish (MECP, 2017), and bw is the body weight of an average healthy adult (assumed 70 kg). We then calculated the margin of exposure (MOE) for microcystins (Eq. 3), which is commonly used in human health risk assessment to assess for potentially genotoxic or carcinogenic compounds.

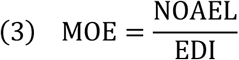

Since the WHO has shown evidence that microcystins are potentially carcinogenic to humans and animals, an MOE ≥ 10,000 was considered for this study to be protective for genotoxic and carcinogenic compounds. The value of 10,000 is derived by the multiplication of two UFs; a 100-fold difference between the calculated reference point and human exposure is applied for species differences and human variability, and an additional 100-fold difference is applied for inter-individual variability (EFSA, 2005). Although it is widely documented that MOEs of 10,000 are used as a protective value in the evaluation of genotoxic and carcinogenic compounds present in food where a single toxicant is being evaluated (EFSA, 2005), the extent to which microcystins is a risk remains unclear for MOEs ≤ 10,000. We therefore adopted the following categories to assist with MOE classification: 1–1,000 (high risk), 1,000–10,000 (medium risk), 10,000–100,000 (low risk) (Cunningham et al., 2011).

We finally calculated the hazard quotient (HQ) for microcystins (Eq. 4) as the ratio of the potential exposure to microcystins and the level at which no adverse effects are expected,

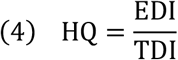

For adverse effects from microcystin exposure, HQ ≤ 0.1 is considered negligible, HQ ≥ 0.1 ≤ 1 is considered low, HQ ≥ 1 ≤ 10 indicates some hazard, and HQ ≥ 10 indicates a high hazard.

### 2.5. Statistical analysis

For statistical analysis any values below the limit of detection of the LC-MS of 2.18 ng g^-1^ ww were treated as zero concentrations. Linear mixed effect models with species treated as a random effect were used to test whether concentrations of microcystins in the muscle and liver were greater than the limit of detection.

## 3. Results and Discussion

Fish muscles contained microcystins (F_1,117_=20.42, *p* <0.0001), with a mean concentration of 1.29 ng g^-1^ ww across species (Figure 1). Only 24/124 samples had concentrations above the limit of detection. 2/124 of samples had concentrations exceeding the WHO’s TDI and the human health risk assessment suggests microcystins are a low risk pre- and post-HAB periods (Table 1). Applying these results against the fish consumption advisory benchmarks in Ontario (Shahmohamadloo *et al*., 2022b), 95% of muscles were <6 ng g^-1^ ww microcystins, which would be classified in the ‘unrestricted’ fish consumption advisory, and all muscles are in the ‘unrestricted’ meal category (<25 ng g^-1^ ww) in Ohio. Considering 98% of muscles had microcystins below the WHO’s TDI, our results indicate that humans can safely consume Lake St. Clair Largemouth Bass, Northern Pike, Smallmouth Bass, Rock Bass, Walleye, White Bass, and Yellow Perch fillets ranging from three times a week to every day (12 to 32 meals per month), depending on the species and its size. A caveat which concerns the validity of our risk assessment result is that these fish were collected pre- and post-HAB periods. A comprehensive review on wild, freshwater fish found variable microcystin concentrations in muscles ranging from 0 to 3.27 μg g^-1^ dw during HABs, yet it was noted that this variability may be due to major limitations in methodology such as the use of ELISA (Flores et al., 2018). More recent work using the Lemieux Oxidation method on Nile Tilapia (*Oreochromis niloticus*) collected from fish ponds before, during, and after HABs found total microcystins (free and bound) could surpass the WHO’s EDI, particularly during the warm months when cyanobacterial blooms are most active (Mohamed et al., 2020). Nile Tilapia are planktivorous fish, and several studies have shown elevated microcystin concentrations in bottom-feeders (Adamovský et al., 2007; Flores et al., 2018; Mohamed et al., 2020; Xie et al., 2004; Zhang et al., 2009). Therefore, caution should be exercised when catching fish before, during, and after HAB events and can depend on species type.

**Figure 1.**
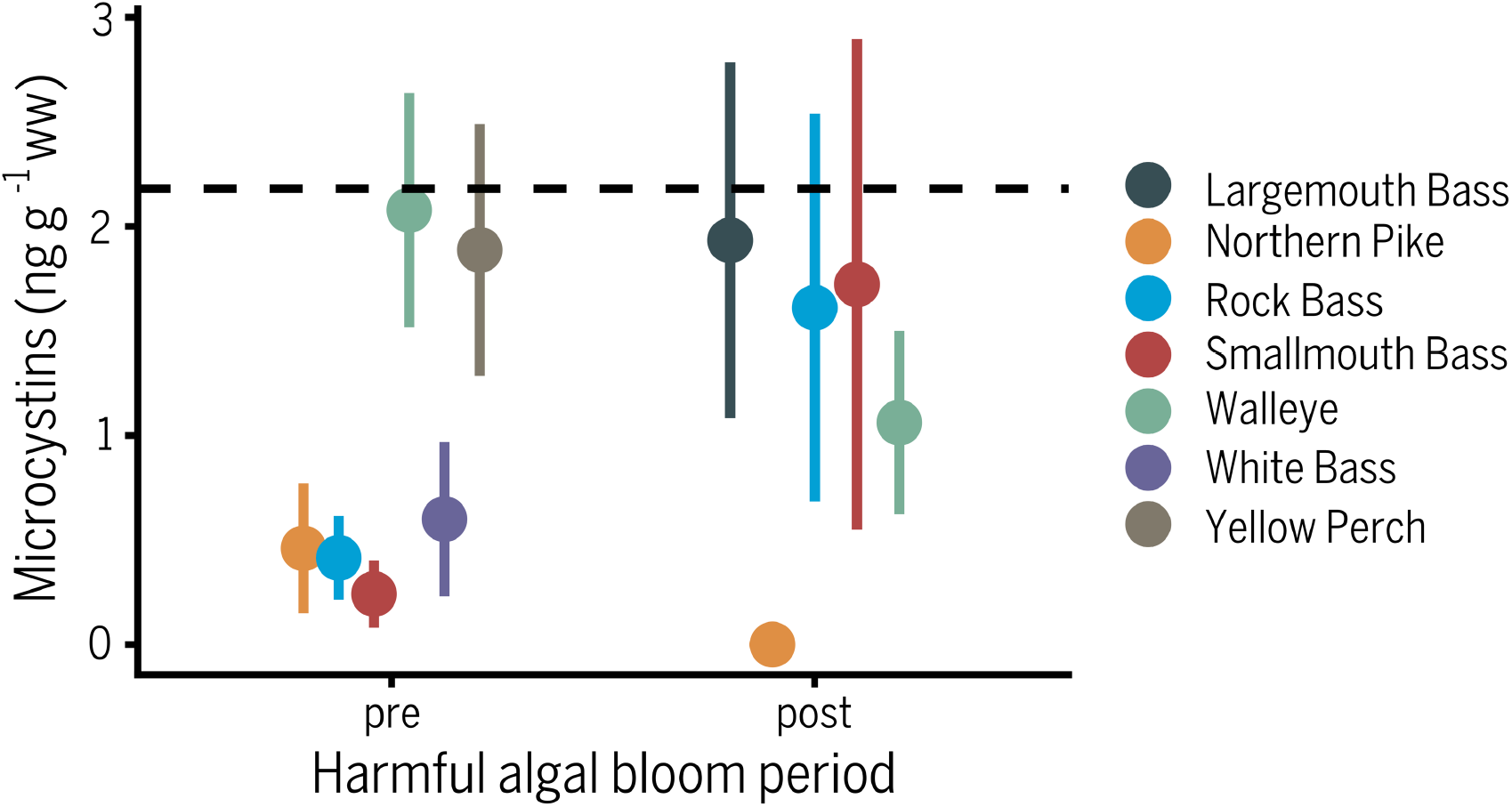
Total microcystins (ng g^-1^ ww) in muscles across seven fish species (Largemouth Bass, Northern Pike, Smallmouth Bass, Rock Bass, Walleye, White Bass, Yellow Perch) collected pre- and post-harmful algal bloom in Lake St. Clair. The method detection limit (2.18 ng g^-1^ ww) is denoted by a dashed line. Refer to Table 1 for the number of fish analyzed per species.

Microcystin concentrations in livers were greater than in muscles with 34/35 samples exceeding the limits of detection. Fish livers contained a significant amount of microcystins (F_1,32_=21.33, *p* <0.0001) with an average value of 224.81 ng g^-1^ ww across Walleye, White Bass, and Yellow Perch (Figure 2; Table 1). The wide variability and presence of microcystin levels in all livers (1-1,500 ng g^-1^ ww) supports our previous work in Lake Erie demonstrating that microcystins are present in fish livers well beyond the cessation of HABs and may be a persistent contaminant in aquatic ecosystems (Shahmohamadloo et al., 2023a). Microcystin-producing HAB species such as *Microcystis*, cyanobacteria common to Lake St. Clair (Davis et al., 2014) and around the globe (Harke et al., 2016), also possess genetic and ecophysiological diversity that allows them to effectively overwinter on the sediments and rapidly form blooms under favorable conditions (Dick et al., 2021). Putative evolutionary adaptations in species such as *Microcystis* (e.g., key genes involved in biosynthesis of secondary metabolites, carbon concentrating mechanisms, detoxification of reactive oxygen species, and acquisition of nitrogen and phosphorus (Dick et al., 2021)) can help to explain the microcystin burden evident in fish livers pre-HAB and offers further insight into the decades-long postulation that microcystins are one among several stress factors involved in fish kills during HAB events (Ibelings et al., 2005).

**Figure 2.**
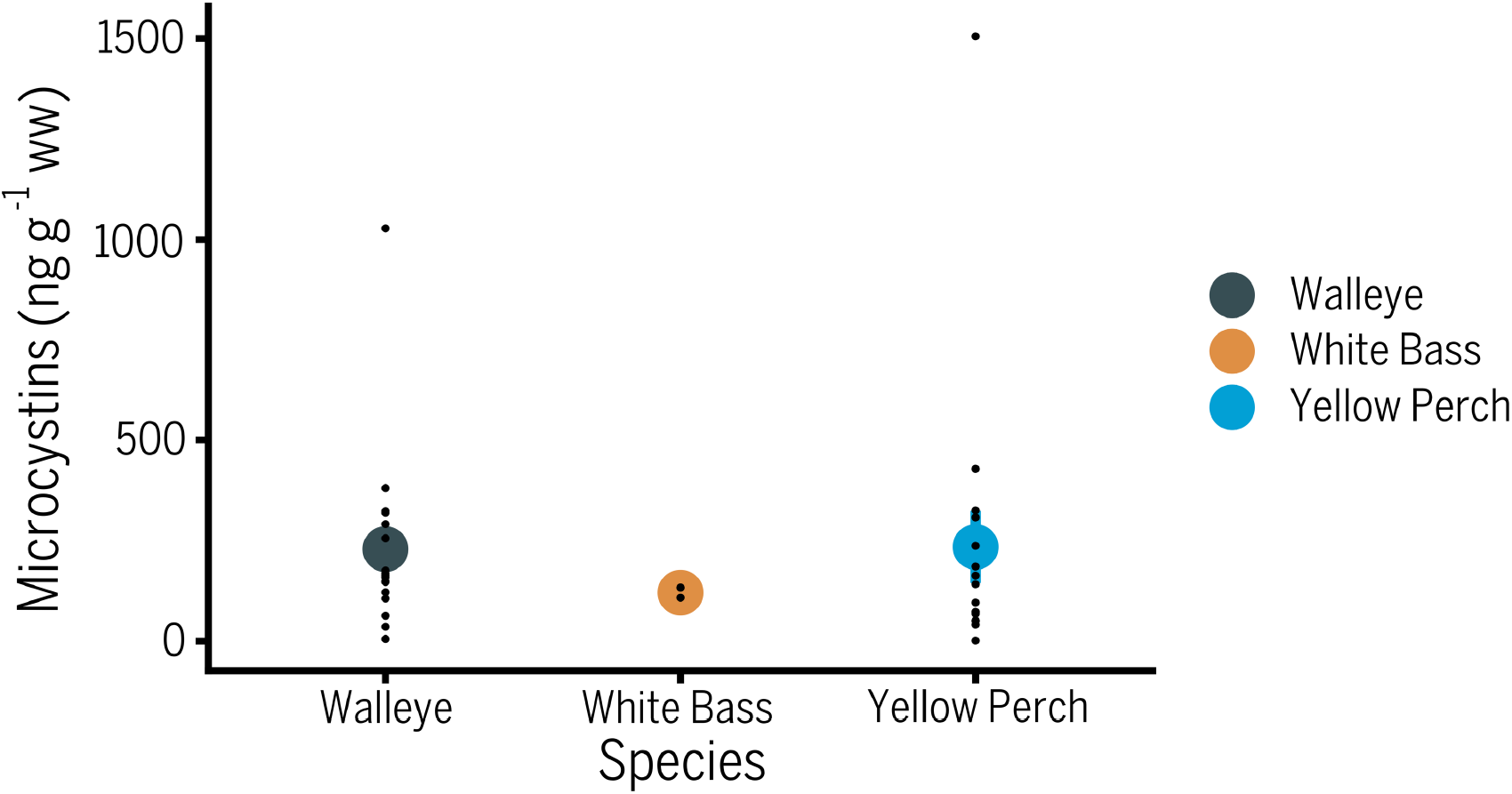
Total microcystins (ng g^-1^ ww) in livers across three fish species (Walleye, White Bass, Yellow Perch) collected pre- and post-harmful algal bloom in Lake St. Clair. The method detection limit (2.18 ng g^-1^ ww) is denoted by a dashed line. Refer to Table 1 for the number of fish analyzed per species.

To alleviate concerns of microcystin exposure via human consumption, general and sensitive populations (women of child-bearing age and children) should avoid eating fish organs from Lake St. Clair and other waterbodies that experience regular HAB events. Our advice is consistent with current advisories for many other contaminants which could be elevated in fish organs. While it is convention to gut fish and strictly consume fillets, whole fish consumption is practiced in various cultures and encouraged for those seeking additional nutrients in their diet (Beveridge et al., 2013; He, 2009). However, microcystins do not break down during cooking and boiling (Harada et al., 1996) and may therefore pose some level of risk to consumers eating fish whole. Considering 90% of anglers consume 8 or less fish meals per month (Awad, 2006) and following the above recommended practices, our results suggest the risk of microcystin toxicity from fish consumption may be low for Lake St. Clair before and after HAB events when fishing is active. Our findings corroborate previous work suggesting microcystin elimination rates from —or degradation rates in— fish tissues can range from days to weeks, depending on a variety of factors including fish species, fish tissue type, and environmental signals (Adamovský et al., 2007; Gurbuz et al., 2016; Shahmohamadloo et al., 2022).

Understanding the potential risks of consuming fish exposed to microcystins, and cyanotoxins more broadly, is critical towards interpreting human and ecological risks. A One Health approach to safeguarding human and animal health offers an interdisciplinary and collaborative path forward in monitoring illnesses or deaths associated with HABs (Hilborn and Beasley, 2015). We therefore exercise prudence in our recommendations and call attention to the need for further work to investigate the chronic and sublethal impacts of HABs and associated toxins on fish populations.

## Acknowledgements

This work was supported by an NSERC CREATE Grant (2013-432269), a Banting Postdoctoral Fellowship, a Canada-Ontario Agreement (2218) through the Ontario Ministry of the Environment, Conservation and Parks, and a Government of Ontario Grant (GLS 1403). Financial support by the Government of Ontario does not equal endorsement of this paper. We thank Ngan Diep for providing microcystin water measurements and the field technicians from the Ontario Ministry of Natural Resources and Forestry for collecting fish and water samples from Lake St. Clair for this study.

